# Model of intracellular ATP production reproduces common electrophysiological signatures of anesthesia

**DOI:** 10.1101/2020.06.17.157149

**Authors:** Pangyu Joo, Heonsoo Lee, Shiyong Wang, Seunghwan Kim, Anthony G. Hudetz

## Abstract

Accumulating evidence suggest that general anesthetics with diverse chemical structure reduce cerebral metabolism with consequent reduction of intracellular adenosine triphosphate (ATP) levels. How cerebral hypometabolism is associated with the typical electroencephalographic (EEG) changes under general anesthesia remains largely unknown.,. We hypothesized that the deficit in ATP production would reduce high-frequency activity, increase low-frequency activity, and cause burst suppression, which are common dose-dependent anesthetic effects on the EEG. To test the hypothesis, we developed a novel neural network model consisting of leaky integrate-and-fire neurons with additional dependency on ATP dynamics. The effect of varying rate of ATP production on neuronal and population activity patterns was simulated under various excitatory/inhibitory balance conditions. A decrease of ATP production suppressed neuronal spiking and enhanced synchronization of neurons over a range of excitatory/inhibitory synaptic strength ratios. As anticipated, the initially asynchronous fast activity was replaced by globally desynchronized slow oscillation and, on further decrease of ATP production, changed into burst suppression with enhanced global synchronization. This study substantiates a novel biophysical mechanism for anesthetic-induced EEG changes through a relationship between energy production and synchronization of neural network.

## Introduction

General anesthesia causes a significant perturbation of neuronal dynamics in the brain. Different kinds of anesthetics have different molecular mechanisms and targets of action (Alkire et al., 2008; Brown et al., 2010; Müller et al., 2011; Purdon et al., 2015) as well as different electrophysiological effects as they produce loss of consciousness. A relatively common electrocortical effect of most inhalational and intravenous general anesthetics with predominantly GABA-ergic action intravenous is an increase of low frequency power of the electroencephalogram (EEG) and local field potential (LFP), frequently accompanied by a decrease in high-frequency gamma band power (Pal, Hudetz, others).. For example, slow oscillation (0.1~1Hz) is commonly observed with the intravenous anesthetics propofol and dexmedetomidine, as well as most inhalational anesthetics (Purdon et al., 2015). An increase of low frequency power can also occur in the delta (1~4Hz), theta (4-8Hz) or alpha (8-14Hz) frequency range, depending on the particular anesthetic used. Further increase in the dose of most anesthetics induces burst suppression pattern in EEG/LFP (Akrawi et al., 1996; Purdon et al., 2015), during which periods of high voltage activity alternate with periods of isoelectricity.

The progressive increase of low frequency EEG activity with deepening anesthesia is thought to be related to a progressive increase in synchronization of neuronal firing. For instance, slow oscillation is characterized by distinct Up and Down states, during which neurons fire together and rest together, respectively, for brief periods lasting in the order of seconds. The enhanced neuronal synchrony facilitates regularized, low-frequency dominant LFP and EEG, leading to a suppression of signal entropy and complexity (Silva et al., 2012; Kreuzer et al., 2014; Liang et al., 2015; Wang et al., 2017). At moderate levels of anesthesia neuronal synchrony may be initially localized such that distant regions show incoherent oscillations (Lewis et al., 2012). As the anesthesia is made deeper, global coherence emerges along with burst suppression (Erchova et al., 2002; Vizuete et al., 2014).

The generally accepted mechanism of anesthetic action in the nervous system involves a primary effect on neurotransmission via synaptic and extrasynaptic action, modulating voltage-gated and ligand-gated ion channels, neurotransmitter release, postsynaptic potentials, neuronal excitability and action potential firing. It is also well established that most general anesthetics significantly reduce cerebral blood flow and cerebral metabolic rate of oxygen (Alkire et al., 1995; Kaisti et al., 2002; Brown et al., 2010; Hudetz, 2012), which are commonly attributed the coupling of cerebral metabolism to neuronal activity. However, the alternative proposal can be made that the anesthetic suppression of cerebral metabolic rate could be, at least in part, to due to a direct inhibition of mitochondrial energy production. In line with this proposal, Kishikawa et al., recently found that three distinct anesthetics, isoflurane, pentobarbital, and 1-phenoxy-2-propanol (a non-clinical agent) could all abolish mitochondrial membrane potential, resulting in an inhibition of mitochondrial adenosine triphosphate (ATP) synthesis (Kishikawa et al., 2018). Consequently, we were interested in the question if metabolic suppression itself may reproduce at least some of the well-known electrophysiological effects of general anesthesia.

In this study we investigated the effect of diminished cerebral metabolism on electrocortical activity using a neuronal network model. The dynamical pattern of neuron firing was explored under different metabolic conditions and at various excitatory/inhibitory synaptic ratios, asking if lowering intracellular ATP production rate would reproduce known characteristics of neuronal network dynamics, neuronal synchrony and frequency of population activity as commonly observed under anesthesia.

## Materials and Methods

### Experimental procedures

The study was approved by the Institutional Animal Care and Use Committee in accordance with the Guide for the Care and Use of Laboratory Animals of the Governing Board of the National Research Council (National Academy Press, Washington, D.C., 2011). A multi electrode array consisting of 64-contact silicon probes (shank length 2mm, width 28-60 μm, probe thickness 15 μm, shank spacing 200 μm, row separation 100 μm, contact size 413 μm2; custom design a 8×8_edge_2mm100_200_413, Neuronexus Technologies, Ann Arbor, MI) was chronically implanted in the primary visual cortex of each animal (eight adult male Long-Evans rat). The tip of the probe was at 1.6 mm below the pial surface. For recording the electromyogram, a pair of insulated wires (PlasticsOne, Inc., Roanoke, VA) exposed at their tip was placed bilaterally into the nuchal muscles.

One to 8 days after surgery, the volatile anesthetic desflurane was administrated in stepwise decreasing concentration at 6%, 4%, 2%, and 0%, one to eight days after surgery. A 15-minute equilibrium period was allowed to stabilize the anesthetic concentration between consecutive conditions. At each anesthetic concentration neuronal activity was recorded first during spontaneous activity for 20 minutes followed a period with visual stimulation (light flashes delivered to the retina by transcranial illumination). Data obtained with visual stimulation were not used in this study. In one experiment that was performed in the beginning of the study, only forty minutes of spontaneous activity was recorded (ten minutes per anesthetic concentration). Because data were analyzed from 10-second epochs, the shorter 10-minute data length in this experiment was considered inconsequential.

Extracellular potentials were recorded at 30 kHz sampling rate (SmartBox, Neuronexus Technologies, Ann Arbor, MI). For spike recording the signals were median-referenced and high-pass filtered (300 Hz). For LFP the signals were bandpass-filtered at 0.1 to 250 Hz. Signals with absolute value greater than 10 SD were considered movement-related artifacts and automatically excluded from the analysis. The data were also visually inspected and noticeable noise episodes were manually excluded. One experiment was excluded from the analysis due to severe noise contamination. A template-based spike sorting method, Spiking Circus (Yger et al., 2018) was used to identify single unit activity (SUA). Per each animal, 36 ± 14 (mean ± SD) single units were obtained.

### Model

#### Model neuron

The ATP-gated potassium channel has been suggested as a key component in the networks of Hodgkin-Huxley type neurons that exhibit metabolism-dependent slow activities (Cunningham et al., 2006; Ching et al., 2012). To efficiently simulate metabolic-dependent activity in a large network, we constructed a simplified neuronal model by adding an ATP-dependent current term that behaves like the ATP-gated potassium channel in leaky integrate and fire neurons. The membrane voltage – current equation is

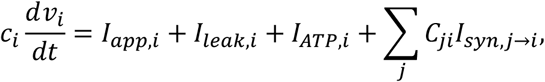

where *I_app_* is an externally driven constant current (*I_app_* = 0.1), *I_leak_* is leakage current with time constant 38.75 ms (Lansky et al., 2006), *I_ATP_* is the ATP-dependent current term, *I_syn_* is the synaptic current, and *C_ij_* is a constant of synaptic strength from yth neuron to *i*th neuron. *v* is defined to range from 0 to 1 and the capacitance *c_i_* is defined as 1. Additionally, each neuron is driven by an independent Poisson input with a mean of 0.1 Hz, which increases the membrane voltage *v* by 0.5. The dynamical equations for *I_ATP_* are

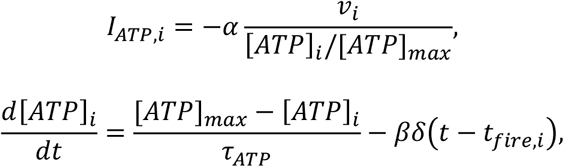

where *α* is the conductance of the ATP-gated potassium channel, *β* is the ATP consumption per each spike and *τ_ATP_* is the time constant for ATP recovery from mitochondrial energy production. The parameter values for *α* = 0.15, *β* = 0.001, [*ATP*]_*max*_ = 1, and *τ_ATP_* varies from 8 sec to 40 sec. 1% (SD) randomness on *I_app_* and *τ_ATP_* are given for each neuron.

For a wide range of gamma-aminobutyric acid (GABA) agonist type anesthetics, cerebral metabolic rate dramatically decreases with deepening anesthesia. Anesthetics affect on the mitochondrial respiratory chain or its structure, such as an inhibition of respiratory complex I or II, with decreased mitochondrial ATP production (La Monaca and Fodale, 2012). Moreover, a recent study demonstrates that the inhibition of mitochondrial ATP synthesis can result from abolished mitochondrial membrane potentials under the effects of anesthetics (Kishikawa et al., 2018). Therefore, in the model we modulated *τ_ATP_*, which is the inverse of the ATP production rate of a single neuron, so that the neurons have a larger *τ_ATP_* at a deeper level of anesthesia.

The numerical simulation was performed in MATLAB with a 0.5 ms time step using a second order Runge-Kutta method. The simulations were run for a time period of 100 sec, and the first 20 sec was removed before analysis to avoid undesired transient effects.

#### Network architecture

We constructed a network of leaky integrate and fire neurons including 8,000 excitatory and 2,000 inhibitory neurons (Fig. 1). The neurons are randomly scattered on 2D rectangular plane (5 mm by 20 mm) and make contacts to the nearby neurons with a probability that depends on the spatial distance between two neurons. The probability distribution follows a Gaussian distribution that is centered at 0 and has different standard deviation σ for excitatory neurons (σ_ex_ = 250 *μm*), and inhibitory neurons (σ_inh_ = 125 *μm*) (Compte et al., 2003), and the resulting node degree of the network is 32 ± 7 (SD).

**Figure 1.**
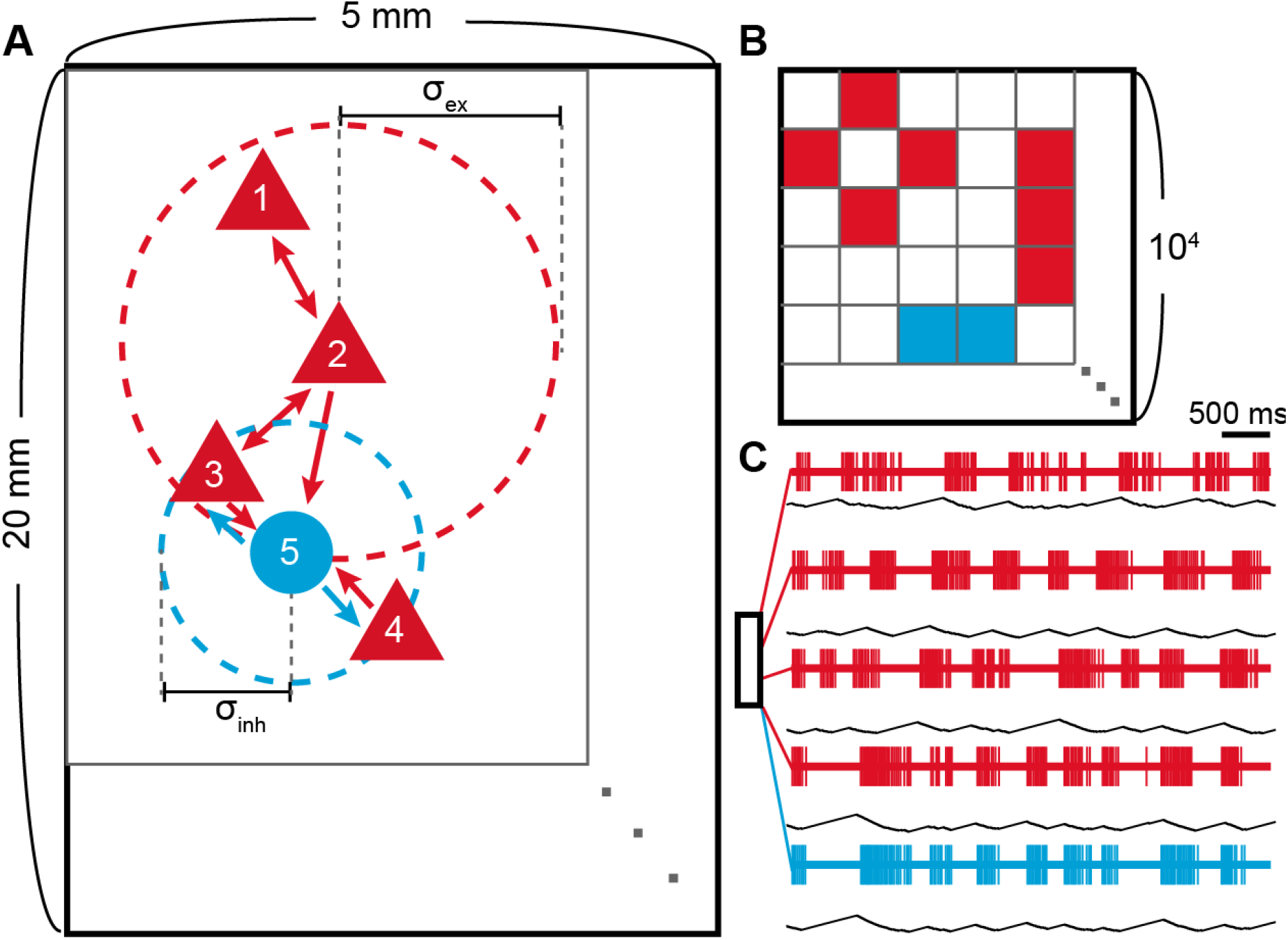
Model schematic and spiking patterns. A) Schematic representation of the model network. The red triangle represents an excitatory neuron, and the blue circle represents an inhibitory neuron. The outer circles with dashed line around neuron 2 and 5 correspond to the connection range of the neuron 2 and 5, respectively. B) Connectivity matrix (*C_ji_*; 10^4^ by 10^4^) of the schematic. Synaptic connection is made from *j* to *i*. C) Location dependent spiking patterns in the model. Red and blue lines represent single neuron spiking activities and the black line below represents corresponding ATP level of the neuron. Small box represents the 20 mm by 5 mm rectangle and the five neurons are selected according to their location. The two bottom neurons are selected from nearby locations and therefore show similar spike patterns.

Incoming synaptic current *I_syn_* induces a perturbation on the membrane voltage *v*. The functional form of the postsynaptic current follows 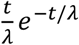, where *λ* is different for the excitatory (*λ_EPSC_*=2 ms) and the inhibitory (*λ_IPSC_* = 5 ms). As inhibitory-to-excitatory (I/E) balance could have a significant effect on the dynamics, we manipulated I/E balance as an additional parameter by multiplying a ratio constant r on the inhibitory *C_ij_* and r varied from 0 to 6. When r = 1, the excitatory *C_ij_* is 0.25 and the inhibitory *C_ij_* is −0.1 so that the cumulative sum of a single postsynaptic current for an excitatory spike and an inhibitory spike are the same. When r = 2, on the other hand, the inhibitory *C_ij_* becomes −0.2 whereas excitatory *C_ij_* remains to be 0.25.

#### Simulated LFP

We calculated simulated local field potentials (sLFP) to characterize collective behavior of the system and to compare it to the experimental data. sLFP is calculated by summing up the excitatory post synaptic currents (EPSC) weighted by a shape function *f*(*l*).

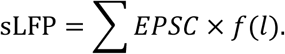

Here, *f*(*l*) is a weighting function that only depends on the distance (*l*) between the measuring point and a neuron; thus, it represents a single neuron contribution on LFP. A detailed simulation study has investigated the form of the shape function (Lindén et al., 2011) and we utilized roughly the distance dependency of the function. In this study, *f*(*l*) is flat when *l* < 100 μm, and follows *l*^−2^ scaling when *l* > 100μm.

## Results

### Anesthesia suppresses total spike activity and promotes synchronization

First we examined how anesthesia alters the amount of spike activity and synchronization in experimental data. Figure 2A depicts raster plot and corresponding LFP signal at each level of anesthesia from a representative animal. The number of spikes gradually decreased from S1 to S4. In addition, the temporal dynamics of population activity was profoundly altered. S1 was characterized by irregular firing pattern while the LFP signal showed low amplitude and high frequency. As the anesthesia deepened, the neuronal spikes became temporally fragmented as indicated by virtually silent periods between bursts of spikes. Simultaneously, the LFP amplitude increased and the frequency shifted to slow oscillation. Burst suppression pattern appeared in deep anesthesia (S4).

**Figure 2.**
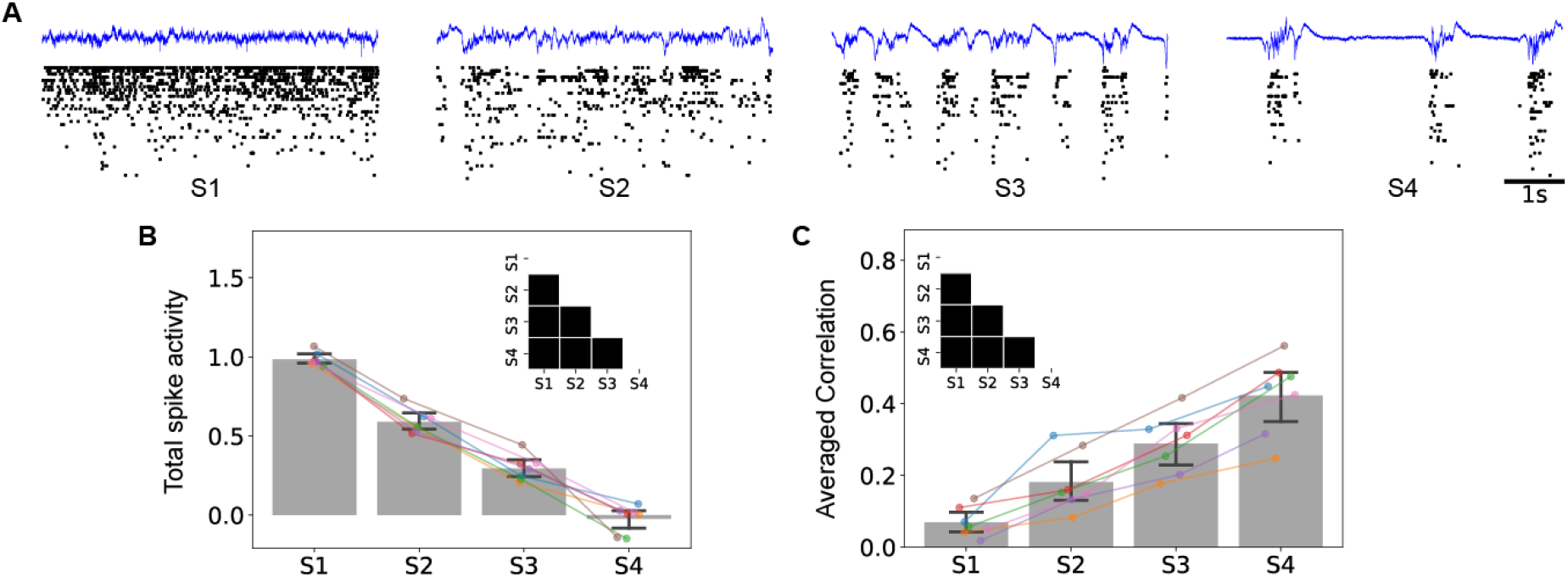
Suppressed spike activity and enhanced synchronization in anesthesia. A) Raster plot (black dots) and LFP trace (blue line) under different depths of anesthesia; S1-S4 corresponds to different depths of anesthesia (from wakefulness to deep anesthesia). Note the increased bursting as anesthesia deepens. B) Total spike activity decreases monotonically. C) Averaged correlation increased monotonically. In B) and C), the error bar indicates 95% confidence interval across seven animals. The inset in each panel represents statistically significant difference between pairs of anesthetic states.

To test whether the finding is observed in all animals, we calculated total spike activity, that is, the total sum of the number of spikes in a neural network, and global synchronization, which is estimated by the averaged pair-wise correlation of all neurons. Because the number of neurons recorded in each experiment was different, the total spike activity at each level of anesthesia was normalized to the total spike activity in wakefulness (S1) of the same animal and then averaged over seven animals. Figure 2B shows that the total spike activity of all animals decreased from S1 to S4. Meanwhile, the averaged correlation increased from S1 to S4 (Fig. 2C). For both total spike activity and averaged correlation, statistical significance was found for all pairwise comparisons between the four levels of anesthesia.

### Neuronal network dynamics at diminished ATP recovery

The overall dynamical patterns were examined with varying ATP recovery time constant, *τ_ATP_*. In addition, because anesthetics alter the balance of excitation and inhibition, we manipulated the E/I balance (r) as an additional parameter. The model showed several distinct firing patterns depending on *r* and *τ_ATP_* (Fig. 3). When the ATP recovery rate was high (*τ_ATP_*=8 sec) the firing rate was high with a constant firing rate over the 20 sec time period. Some neurons had low firing rates (white horizontal lines in Fig. 3) due to lack of excitatory connections or excess inhibitory connections. As *τ_ATP_* decreased to 12 sec, the firing rate decreased without oscillatory pattern. At *τ_ATP_*=14 sec, slow patterns near 1 Hz began to appear, and the oscillation slowed down as *τ_ATP_* was lengthened. As was lengthened further, the spiking pattern acquired alternating burst and silent phases, which was similar to the burst suppression in deep anesthesia.

**Figure 3.**
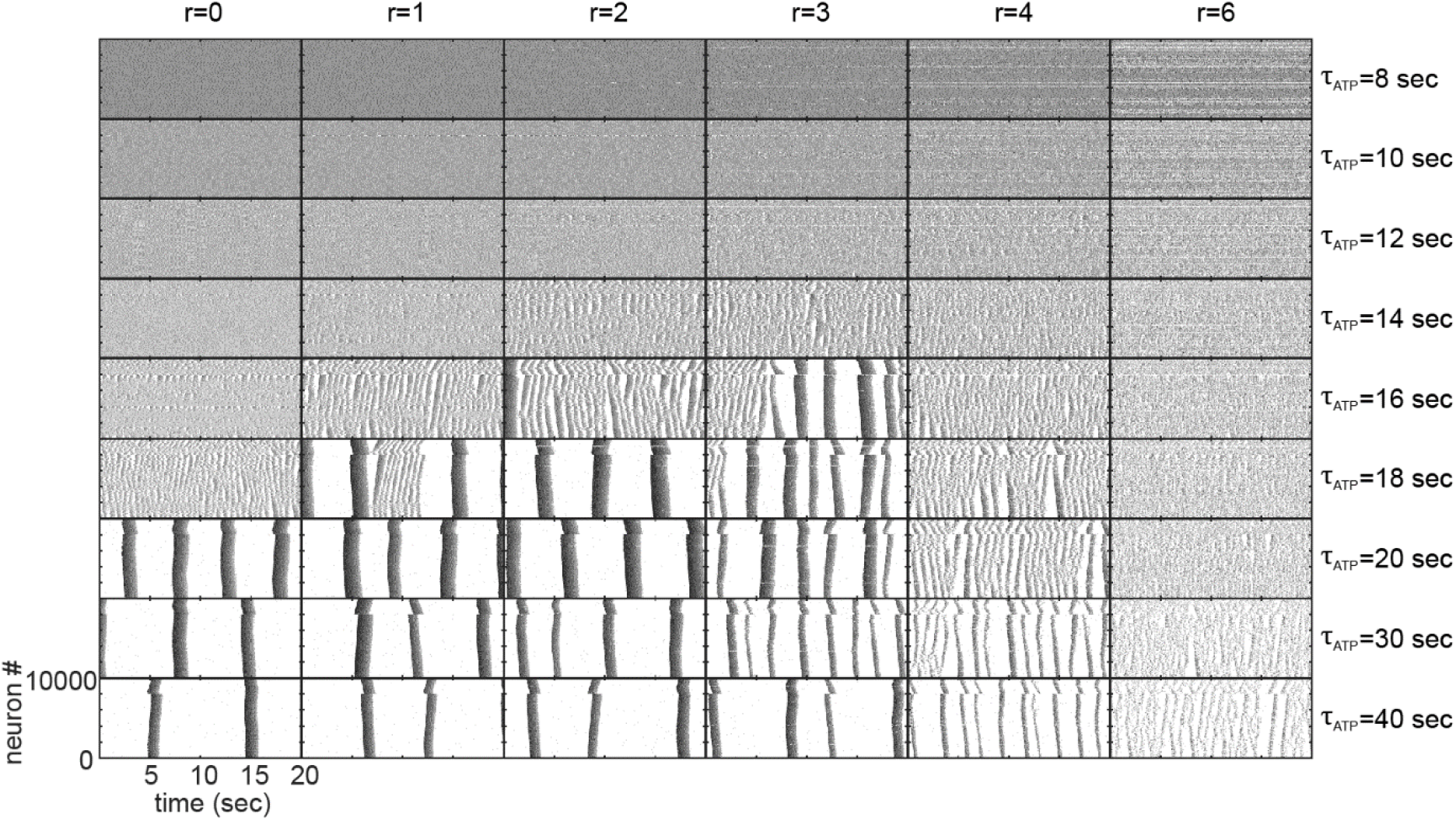
Spike activity at different I/E ratios (r) and ATP recovery rate time constant. Higher firing rate is indicated by darker shade. In each box, the order of 10^4^ neurons on the y axis is determined by their type, excitatory (1 to 8,000) or inhibitory (8,001 to 10,000), and location (20 mm side). The last 20 sec of the simulation is shown.

Interestingly, the different I/E balance cases exhibited qualitatively similar pattern changes associated with *τ_ATP_*; the stronger inhibition led to more irregularities and shorter durations for collective firing, but it did not affect much on the qualitative patterns of the emergent synchrony (Fig. 3).

To illustrate the predictive power of the model, we manually selected from all simulation results four states (S1’-S4’) that visually corresponded to the spiking activity S1-S4 in our experiment. A selected state with a higher number corresponded to a deeper level of anesthesia with longer *τ_ATP_* and thus slower ATP recovery with the slowest being S4’ (*τ_ATP_* = 8, 12, 16, and 40 sec for S1’, S2’, S3’, and S4’, respectively with r = 3). S1’ exhibited fast and irregular firing patterns with the highest firing rate of all states. From S1’ through S2’ to S3, oscillatory pattern appeared with increasing amplitude in sLFP. At S4’, the network exhibited bursting periods, which have very high spiking rate (up to 80 Hz), and silent periods, which showed near zero firing rate that lasted for several seconds. The total spike activity was calculated from the simulated spikes and normalized with S1’. As with the experimental data, the total spike activity decreased from S1’ to S4’ (Fig. 4B). The average spike correlation of all pairs of neurons (C_tot_) and of neuron pairs located within 200 *μm* (C_local_) both monotonically increased from S1’ to S4’ (Fig. 4.C).

**Figure 4.**
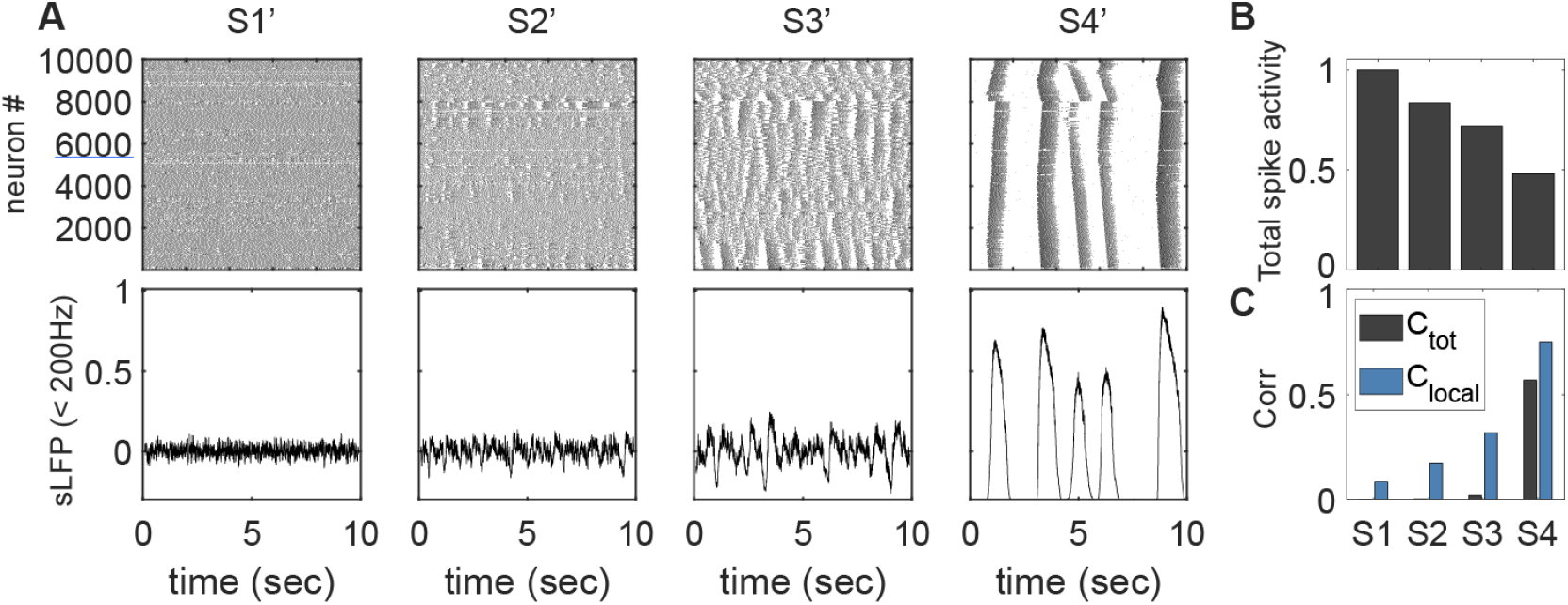
Quantitative characteristics and simulated LFP of four selected states. A) Raster plot (top) and simulated LFP (bottom). The last 10 sec of S1’ (*τ_ATP_* = 8 sec), S2’ (*τ_ATP_* = 12 sec), S3’ (*τ_ATP_* = 16 sec), and S4’ (*τ_ATP_* = 40 sec) are visualized. B) Total spike activity normalized to total spike activity in S1’. C) Average pairwise correlation of all neurons in the network (*C*_tot_) and of nearby neurons within 200 *μm* (*C*_local_). The total spike activity and averaged correlation are calculated from 80 sec of the simulation.

### Phase diagram analysis

To illuminate the mechanism of neuronal interactions underlying the state transitions, we performed a phase plane analysis (Fig. 5.). We chose mean firing rate (MFR) and intracellular concentration of ATP ([ATP]) as the representative coordinates. MFR and [ATP] were averaged over 200 ms moving windows with 50 ms time step for a total 10^4^ neurons (black arrows in Fig. 5B) or local 100 neurons at the center of the square plane (blue arrows in Fig. 5B). Up and Down periods were defined using the temporal MFR of an neuron with 20 Hz threshold (Fig. 5C).

**Figure 5.**
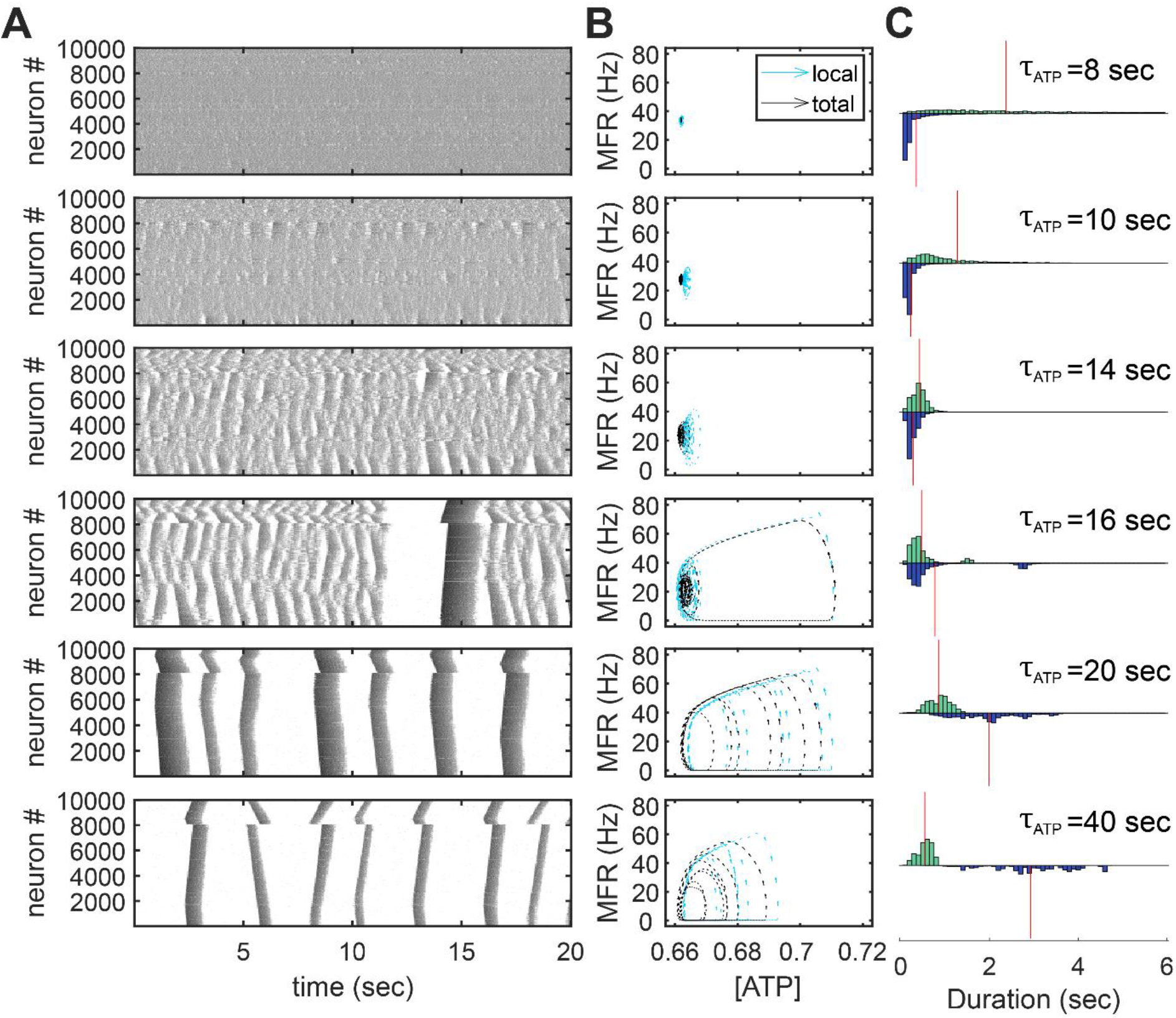
State transitions of neuronal network dynamics (r=3). A) Raster plots. The graded shades of grey represent the firing rate; darker for higher and lighter for lower firing rate. Each row of panels correspond to a different ATP recovery time constant (from top to bottom, *τ_ATP_* = 8, 10, 14, 16, 20, 40 sec, respectively). B) Phase diagrams. The arrows represent positions and velocities on the phase plane, which are averaged over 200 ms moving window with 10^2^ (local) or 10^4^ (total) neurons. C) Duration of Up and Down states. The upper green bar and the lower blue bar represent histogram of Up state duration and Down state duration respectively; and the red vertical line marks the average value.

The emergence mechanism of the slow oscillation in our model is based on the excitatory feedback from the network and the slow intracellular ATP dynamics as suggested by (Cunningham et al., 2006). When the ATP recovery is sufficiently high (*τ_ATP_* = 8 sec, first row in Fig. 5), the firing rate is relatively stable and without any silent period. As the ATP recovery rate starts to decrease (*τ_ATP_* = 10 sec), the reduced ATP level results in a suppression of firing rate. At *τ_ATP_*= 14 sec, the reduced ATP recovery rate does not only reduce firing rate, but also profoundly alters temporal dynamics of neural network activity; at this stage, both MFR and [ATP] become locally synchronous (third row in Fig. 5B). Because a spike from an excitatory neuron can trigger spikes in the other neurons, which in turn, promotes the next spike of the same neuron (i.e., excitatory feedback), the neural network enters to a transiently high activity pattern (Up state) followed by a silent period (Down state) (*τ_ATP_*=14 sec, third row in Fig. 5).

At a higher *τ_ATP_* (*τ_ATP_* = 16 sec), a long silence period emerges and is followed by collective bursting of all the neurons firing in synchrony (burst suppression pattern; near 10-15 sec, fourth row in Fig. 5A). The emergence of the burst suppression cycles on the phase plane could be explained by the competition between the propagation speed of synchronous spikes and the resulting suppression (silence) time. For example, at *τ_ATP_* = 14 sec (third row in Fig. 5), globally desynchronized slow oscillations arise from the limited propagation speed of synchronous spikes and then the spikes form a continuous wave without a global silence over the space; a new wave appears before the prior wave fades away. As the ATP recovery slows further (*τ_ATP_* = 16 sec, fourth row in Fig. 5), the length of the silent periods become longer. If the duration of the silent period becomes comparable to the time required for the synchronous spikes to propagate over the whole system, the system cannot sustain the continuous wave because the it fails to have enough activity required to initiate a local Up state (near 10 sec in *τ_ATP_*=16 sec conditions, fourth row in Fig. 5A). In consequence, the system occasionally fall into a globally silent period. This silence persists for a period of time until a sufficiently large number of nearby neurons are able to fire together and initiate synchronous spikes. Notice that ATP level at the end of the silent period at *τ_ATP_*=16 sec is even higher than the low *τ_ATP_* conditions, and is synchronous among neurons (Fig. 5B; e.g., *τ_ATP_* ≤ 14 sec). This leads to burst spikes with a high intensity until they use up the accumulated ATP (near 14 sec in *τ_ATP_*=16 sec conditions, fourth row in Fig. 5A). If the lengths of the inactive periods exceed the time required for a burst to propagate over the whole system, the large burst cycle becomes the only stable dynamical pattern in the phase plane (*τ_ATP_* = 20 sec, fifth row in Fig. 5). When the ATP recovery slows down further, the size of the burst cycle shrinks with lengthened period (*τ_ATP_* = 40 sec, sixth row in Fig. 5) and it finally disappears for 0 recovery rate.

## Discussion

The goal of this study was to test the hypothesis that a change in metabolism-dependent neuronal interactions could account for anesthetic-induced cortical state transitions in a spiking neuron network model. At various levels of excitatory-inhibitory synaptic strength, or E/I balance, the neuronal network showed irregular and fast firing pattern when the ATP recovery rate was high. The firing pattern reorganized into locally synchronous but globally desynchronized slow oscillatory activity at an intermediate ATP recovery rate and into globally synchronous burst firing, or UP/Down states, when ATP recovery rate was further decreased. These results were consistent with experimental data that showed low firing rate and increased synchronization under general anesthesia. The state transitions were then explained by the implicated burst-preference of leaky integrate-and-fire neurons at lower firing rate and the rebound firing after synchronized silence.

### Reduced ATP production alters neural network dynamics

Metabolism has crucial effect on neuronal dynamics in the brain. The brain consumes most of its energy on neuronal activities such as synaptic transmission, ion pumping to maintain the resting potential and generating action potentials (Harris et al., 2012). Accordingly, the firing rate of neurons is highly correlated with the concurrent cerebral metabolic rate (Smith et al., 2002; Mäkiranta et al., 2005; Ekstrom, 2010). The degree of neuronal activity can be regulated by the ATP-gated potassium channel that directly affects the membrane potential as a function of intracellular ATP concentration (Yamada and Inagaki, 2002; Huang et al., 2007; Sun and Feng, 2013). The ATP-gated potassium channel has been included in neuronal models designed to explain the mechanisms of slow oscillation (Cunningham et al., 2006) and anesthetic-induced burst suppression (Ching et al., 2012) to control neuronal spiking as a function of metabolic state.

Anesthetics may influence cerebral neuronal activity directly, through receptor-mediated and biophysical mechanisms, as well as by limiting intracellular high-energy phosphates due to the suppression of mitochondrial respiration. Studies using positron emission tomography revealed that whole brain metabolism is substantially diminished during the administration of propofol or sevoflurane (Alkire et al., 1995; Kaisti et al., 2002). The metabolic suppression is correlated with simultaneous changes in quantitative EEG descriptors (Bispectral Index, total power, relative beta power, etc.) (Alkire, 1998). A causal link between metabolic and electrophysiological activities could be the abolished ATP production by anesthetics. Several commonly used anesthetics directly influence mitochondrial enzymes and metabolic pathways reducing the production of ATP (La Monaca and Fodale, 2012). Abolished mitochondrial membrane potential under isoflurane, pentobarbital, or 1 - phenoxy-2-propanol anesthesia can also inhibit mitochondrial ATP synthesis (Kishikawa et al., 2018).

In line with these studies, our present model study shows that ATP production rate could be a key regulator of the state transitions between irregular wake-like firing, slow oscillation and burst suppression. Our findings, together with the above-cited studies, suggest that the suppression of neuronal activities due to diminished metabolism could be a core principle for the state transitions in general anesthesia.

### Transition from slow oscillation to burst suppression

Burst suppression is a prevalent phenomenon of deep anesthesia and is characterized by alternating burst and suppression periods. Although many studies have been conducted to explain the characteristics of burst suppression, the biophysical mechanism of the burst suppression remains unclear. Based on our model predictions, we suggest that a competition between the propagation speed of synchronous spikes and the length of the silent period may lead to the intermittent transition between slow oscillation and burst suppression. Additionally, we can make some predictions on the intermediate state between slow oscillation and burst suppression. First, the distribution of Down state duration may follow a bimodal distribution. Second, sporadic large fluctuations may be observed before burst suppression with increasing probability as the anesthesia level deepens.

Moreover, we could observe the spatial patterns of the slow activities with our model. Burst suppression has been known to be a predominantly synchronous phenomenon (Clark and Rosner, 1973; Lewis et al., 2013). On the other hand, recent experimental studies have shown that the anesthetic-induced slow oscillations occur asynchronously across cortex (Lewis et al., 2012; Flores et al., 2017). Our model suggests an explanation for the difference in synchronization. In our model, the two states exhibit qualitatively different dynamical patterns. Slow oscillations in EEG??? Spike??? appear globally asynchronous and continuous waves due to their limited propagation speed. On the other hand, during burst suppression, long-lasting silence aligns the individual neurons enabling high level of global synchronization. In this way, the suppression of neuronal activity caused by diminished ATP recovery can lead to the enhanced synchronization without any modification on the network connectivity.

### Hyperexcitability in deep anesthesia

A modeling study about the propagation of slow oscillatory activity on a cortical network suggested that the wave propagation speed is dramatically increased with blockage of inhibition (Compte et al., 2003). In deep anesthesia accompanied burst suppression is characterized by paradoxical hyperexcitability to sensory stimuli (Hudetz et al, 2007), presumably due to diminished inhibition (Kroeger and Amzica, 2007; Ferron et al., 2009). Likewise, in our model, burst suppression occurred at higher ATP recovery rates with lower inhibitory activities (Fig. 2, r<4). Therefore, our model suggests that the two core causes of the burst suppression are the attenuation of overall neuronal activity due to low intracellular ATP and the acceleration of wave propagation (or increase in synchronization) due to disinhibition.

### Cortical model of slow oscillation

Slow oscillation in general anesthesia appears with significantly increased power broadly across temporal and parietal regions and the slow oscillation power increases in a dose dependent manner (Purdon et al., 2013). The slow oscillation has been considered as a mostly cortical phenomenon as shown by its survival after thalamic lesions (Steriade et al., 1993) and many in vitro experiments that have shown slow oscillations with different cortical slices (Neske, 2016). Also, the studies with a model of cortical network that exhibits slow oscillations (Compte et al., 2003; Cunningham et al., 2006) supports the hypothetical origin of the anesthetic slow oscillation. Consistent with previous studies, our model could also provide a cortical mechanism of slow oscillation. Our model showed that the suppression of overall firing rate due to diminished ATP production can cause the occurrence and growth of the slow oscillation. Furthermore, we could observe that the emergence of the slow oscillation starts with occasional occurrences of a brief down state and it morphs into distinct up states and down states with decreased intracellular ATP production.

### Limitations

First, the firing rate in our model network was uniformly distributed across neurons, distinct from many experimental studies, in which firing rate distribution follows log-normal distribution (Buzsáki and Mizuseki, 2014). The uniformity in our model is originated from the homogeneous degree distribution of the lattice like model network. In this sense, our model might represent only a small portion of neurons with many and strong synaptic connections. However, because the synchronization of a highly inhomogeneous neuronal network is dominated by small subset of high degree nodes (Grinstein and Linsker, 2005), the overall dynamics would not be dramatically changed although additional neurons with less and weak synaptic connections were taken into account. Second, we did not reproduce the alpha and beta oscillations, which are associated with sedative and paradoxical excitation states under anesthesia. The alpha and beta oscillations were previously reproduced with a thalamocortical circuit and synaptic modification mechanisms (e.g., GABA agonist effect) (Ching et al., 2010; Hindriks and van Putten, 2012, 2013; Ching and Brown, 2014). Importantly, neural activity in this frequency range depends on the type of anesthetic; e.g., propofol and dexmedetomidine show distinct EEG patterns in the alpha-beta range but similar low-frequency activity in the delta band (Purdon et al., 2015). Thus, neural activities in the alpha-beta frequency range may be related to specific agent and dose-dependent mechanisms and may not be explained solely by the suppression of metabolism.

### Conclusions

The present study proposes a model in which diminished cerebral metabolism plays a key role in anesthetic-induced state transitions as indicated by the predicted EEG/LFP changes. Specifically, our model could explain transitions from irregular firing state through locally synchronized state to globally synchronized state with monotonically decreasing firing rate consistent with the experimental results. The model provides a framework to further investigate biophysical mechanism underlying metabolism-dependent state transitions under anesthesia.

## Acknowledgements

Research reported in this publication was supported by the National Institute of General Medical Sciences of the National Institutes of Health under award number R01-GM056398 and the Center for Consciousness Science, Department of Anesthesiology, University of Michigan Medical School, Ann Arbor, Michigan, USA. The content is solely the responsibility of the authors and does not necessarily represent the official views of the National Institutes of Health. The authors thank Minkyung Kim for valuable comments and discussions.

